# Eliciting OTUD3/RIPK-Dependent Necroptosis to Prevent Epithelial Ovarian Cancer

**DOI:** 10.1101/2020.04.29.069021

**Authors:** Joshua Johnson, Elise C. Bales, Benjamin G. Bitler, Zachary L. Watson

**Affiliations:** Division of Reproductive Sciences, Department of Obstetrics & Gynecology, University of Colorado School of Medicine, Aurora, CO 80045, USA

**Keywords:** epithelial ovarian cancer, HGSOC, prevention, necroptosis, OTUD3, RIPK, progesterone, TNFα

## Abstract

**Background:** There is an urgent need for early prevention strategies against high grade serous ovarian carcinoma (HGSOC), the deadliest gynecologic malignancy. Transformed p53-null fallopian tube epithelium (FTE) cells are precursors of HGSOC that may be eliminated by inducing necroptosis, a programmed form of inflammatory cell death. Induction of necroptosis is dependent upon activation of receptor-interacting serine/threonine-protein kinases 1 and 3 (RIPK1/3). TNFα and progesterone (P4) effectively promote necroptosis. In this study, we explore the activation of necroptosis as an approach to inhibit HGSOC progression.

**Methods:** Using gene ontology sets as a reference, we analyzed publicly available datasets of HGSOC to correlate the expression of necroptosis effectors to clinical outcomes. Using *in vitro* models of HGSOC we evaluated the effect of TNFα, P4, and α-eleostearic acid on necroptosis. In parallel, the necroptosis inhibitor Necrostatin-1 was used to confirm necroptosis-specific cell death.

**Results:** Expression of the P4 receptor (*PGR*) was sharply reduced in a HGSOC cohort compared to normal, nonmalignant FTE. However, several genes involved in necroptosis signaling were elevated in HGSOC, including *TNF* and *RIPK1*. Increased expression of *PGR*, the necroptosis effectors *TNF* and *RIPK1/3*, as well as ovarian tumor domain-containing deubiquitinase 3 (*OTUD3*) were associated with higher overall survival in 484 HGSOC cases. HGSOC cells activated necroptosis in response to P4, TNFα, and α-eleostearic acid treatment, while P4 or TNFα treatment of HGSOC cells increased *TNF*, *RIPK1*, and *OTUD3* expression. OTUD3 is a putative tumor suppressor that stabilizes PTEN and is hypothesized to be functionally similar to the necroptosis inducer, OTUD7B. shRNA knockdown of *OTUD3* resulted in decreased PTEN protein and RIPK1 protein.

**Conclusions:** We conclude that necroptosis activation may be a viable prevention strategy that leads to the elimination of transformed FTE “founder” cells and prevents HGSOC tumorigenesis. Our data indicate that HGSOC cells activate necroptosis in response to P4, TNFα, and α-eleostearic acid, suggesting that established HGSOC cells may also be eliminated by activating necroptosis.

## Background

High grade serous epithelial ovarian cancer (HGSOC) is proposed to originate from transformed secretory fallopian tube epithelial (FTE) cells located on the fimbriated end of the fallopian tube [1–5]. During ovulation, normal FTE cells are exposed to follicular fluid which includes a milieu of inflammatory cytokines [6, 7] and reactive oxygen species (ROS) [8]. This highly mutagenic environment can promote DNA damage and genetic instability [9, 10]. Mutation or loss of p53 in FTE cells is thought to be the first step in transformation given that p53 is mutated or lost in nearly 100% of HGSOC [11]. Strikingly, HGSOC incidence accelerates at the time of menopause. Menopause refers to the cessation of menstrual cyclicity that most often occurs between the ages of 45 and 55 [12]. Consequences of menopause include cessation of ovarian follicle development, loss of ovulation, and the cessation of the development of *corpora lutea* concomitant with a dramatic decrease in hormones including estrogen and progesterone (P4). Several studies have examined either perturbation or activation of the P4 axis in patients with HGSOC [13–15]. In a small phase II study of 33 patients with advanced HGSOC (stage III/IV), high-dose megestrol (synthetic P4) failed to induce an objective anti-tumor response [14]. In contrast, in a large multicenter case control study of 330 patients with HGSOC and 982 matched controls, depot medroxyprogesterone acetate (DMPA) significantly reduced risk of HGSOC and had a significant clinical benefit in 20-40% of pre-, peri-, and postmenopausal ovarian cancer patients [15]. DMPA has a good toxicity profile and is FDA-approved for birth control and treatment of post-menopausal symptoms.

P4 treatment commonly results in the upregulation of a wide array of genes including Tumor Necrosis Factor alpha (TNFα), a factor that can promote programmed cell death (PCD) mechanisms including apoptosis and necroptosis [16]. Necroptosis is induced by TNFα interaction with the TNFα receptor (TNFR) promoting multiple downstream events including the formation of a complex containing TRAF2, cIAP1, TRADD, and the serine/threonine kinase receptor-interacting protein 1 (RIPK1) [17]. The formation of this complex can lead to cell death *via* apoptosis (caspase 8-dependent) or necroptosis (caspase 8-independent) [18], the latter resulting from mixed lineage kinase domain-like protein (MLKL) activation and subsequent mitochondrial dysfunction and ROS production [16]. While a complete mechanistic understanding behind RIPK1-dependent PCD remains unclear, it is hypothesized to depend on post-translational modifications. For example, Ovarian TUmor domain containing Deubiquitinase 7B (OTUD7B) deubiquitination of RIPK1 promotes its cytosolic localization, resulting in RIPK1-RIPK3 interaction and necrotic cell death [19, 20]. The loss of P4 that occurs during menopause could therefore result in decreased TNFα production and diminished necroptosis, allowing cancerous progression of transformed fallopian tube epithelial (FTE) cells.

Mutations in breast cancer susceptibility genes 1 and 2 (BRCA1/2) increase the risk of developing HGSOC by 10-fold [21, 22]. BRCA1/2 carriers undergo early onset or premature menopause more often than non-carriers [23], and when this occurs, exposure to decreased P4 levels begin earlier. It is possible that the earlier loss of ovarian function and decreased P4 levels further increase HGSOC risk. Consistent with this, Wu et al. [16] recently showed that P4 signaling induces necroptosis and clearance of p53-mutated FTE cells. Transformed FTE cells with intact P4 signaling are therefore responsive to signals that induce necroptosis. However, loss of progesterone receptor (PGR) occurs in 75% of HGSOCs, and loss of PGR conveys a worse overall patient survival [24]. The question of whether necroptosis can be induced in established HGSOC by bypassing the P4-PGR axis has not previously been addressed. This study examines necroptosis induction in HGSOC cells through P4-dependent and -independent mechanisms and explores the hypothesis that induction of necroptosis can reduce the survival of both transformed FTE cells and established HGSOC cells.

## Results

### Expression of necroptosis machinery correlates positively with HGSOC patient survival

Using gene ontology (GO) IDs as a reference, we systematically compared mRNA expression between normal, nonmalignant FTE (n=24) and HGSOC samples (n=11) in a publicly available cohort (GSE10971). We analyzed *BRCA1*, *BRCA2*, necroptosis (GO ID 0070266), progesterone receptor signaling (GO ID 0050847), TNF signaling (GO ID 0010803), and TNF receptors. Genes with significantly changed expression between FTE and HGSOC groups are shown in **Table 1**. Notably, progesterone receptor (*PGR*) is significantly downregulated in HGSOC, suggesting that transformed HGSOC cells may exhibit reduced responsiveness to necroptosis signaling by P4, as has been shown for endometrial cancer [25] and ovarian cancer [13]. However, other genes involved in TNF signaling and the necroptotic pathway are upregulated, including *TNF*, *RIPK1*, and *TNFRSF21* **(Fig. 1A)**, indicating that necroptosis may be activated in HGSOC by bypassing the P4-PGR signaling axis. We queried a cohort of 488 ovarian cancers in The Cancer Genome Atlas [11] for mRNA expression of necroptosis pathway genes *OTUD3*, *PGR*, *RIPK1/3*, and *TNF*, and then used KMplot [26] to compare overall survival of patients that expressed High or Low levels of these genes **(Fig. 1B)**. Higher expression of necroptosis machinery correlated positively with overall survival, supporting our hypothesis that elevated necroptosis may be associated with increased tumor cell death.

**Table 1.** Differentially Regulated Gene Expression in Necroptosis, PGR Signaling, and TNF Signaling and Receptors between Nonmalignant FTE and HGSOC.

| Gene Symbol | Probe ID | Log2 FC<sup>#</sup> | Adj. P-value<sup>*</sup> |
| --- | --- | --- | --- |
| BRCA2 | 214727_at | 1.472 | 0.0002 |
| **Necroptosis** |  |  |  |
| TNF | 207113_s_at | 1.544 | 2.45e-06 |
| RBCK1 | 221827_at | 1.499 | 0.00045 |
| PPIF | 201490_s_at | 1.214 | 2.75e-06 |
| MAP3K7 | 206854_s_at | 1.112 | 4.94e-06 |
| PYGL | 202990_at | 1.021 | 0.00334 |
| CYLD | 221903_s_at | 0.983 | 0.00123 |
| TP53 | 211300_s_at | 0.920 | 0.00255 |
| RIPK1 | 226551_at | 0.845 | 0.00063 |
| DNM1L | 203105_s_at | 0.818 | 0.00033 |
| FADD | 202535_at | 0.781 | 0.00002 |
| SPATA2 | 204433_s_at | 0.603 | 0.00478 |
| OTUD6B | 222825_at | 0.497 | 0.04980 |
| IPMK | 239878_at | 0.434 | 0.00924 |
| ITPK1 | 217710_x_at | 0.301 | 0.04730 |
| OTUD7B | 238994_at | -0.497 | 0.04010 |
| PELI1 | 232304_at | -0.736 | 0.00805 |
| FAS | 204781_s_at | -0.780 | 0.04390 |
| YBX3 | 213319_s_at | -1.084 | 0.00441 |
| CFLAR | 235427_at | -1.143 | 2.07e-05 |
| CAV1 | 212097_at | -1.144 | 0.01280 |
| SLC25A4 | 214821_at | -1.176 | 0.01100 |
| **PGR Signaling** |  |  |  |
| PHB | 200658_s_at | 1.343 | 2.54e-06 |
| YAP1 | 213342_at | 1.129 | 0.00022 |
| UBR5 | 208884_s_at | 0.852 | 7.16e-07 |
| SRC | 213324_at | 0.335 | 0.04420 |
| UBE3A | 214980_at | -0.601 | 0.01480 |
| WBP2 | 209117_at | -0.633 | 9.89e-05 |
| PGR | 228554_at | -5.545 | 2.27e-10 |
| **TNF Receptors** |  |  |  |
| TNFRSF21 | 214581_x_at | 2.005 | 4.73e-05 |
| TNFRSF1B | 203508_at | 1.082 | 0.01780 |
| TNFRSF17 | 206641_at | 0.943 | 0.00123 |
| TNFRSF12A | 218368_s_at | 0.851 | 0.02640 |
| TNFRSF9 | 211786_at | 0.803 | 1.80e-05 |
| TNFRSF10B | 210405_x_at | 0.615 | 0.00065 |
| TNFRSF25 | 211841_s_at | 0.465 | 0.01430 |
| TNFRSF19 | 227812_at | -2.815 | 7.73e-07 |
| **TNF Signaling** |  |  |  |
| PELI3 | 235431_s_at | 0.858 | 0.02690 |
| SHARPIN | 220973_s_at | 0.726 | 0.00104 |
| OTULIN | 228382_at | 0.589 | 0.00032 |
| CPNE1 | 206918_s_at | 0.570 | 0.00130 |
| SYK | 209269_s_at | 0.523 | 0.03580 |
| PTPN2 | 213136_at | -0.554 | 0.01980 |
| NOL3 | 221567_at | -0.812 | 0.00421 |
| HIPK1 | 212293_at | -1.040 | 1.07e-07 |
| GAS6 | 1598_g_at | -1.180 | 4.80e-05 |

<sup>#</sup> Log2 fold change of HGSOC (n=11) compared to FTE (n=24)
<sup>\*</sup> P-value corrected for false-discovery rate by Benjamini-Hochberg method

**Fig. 1.**
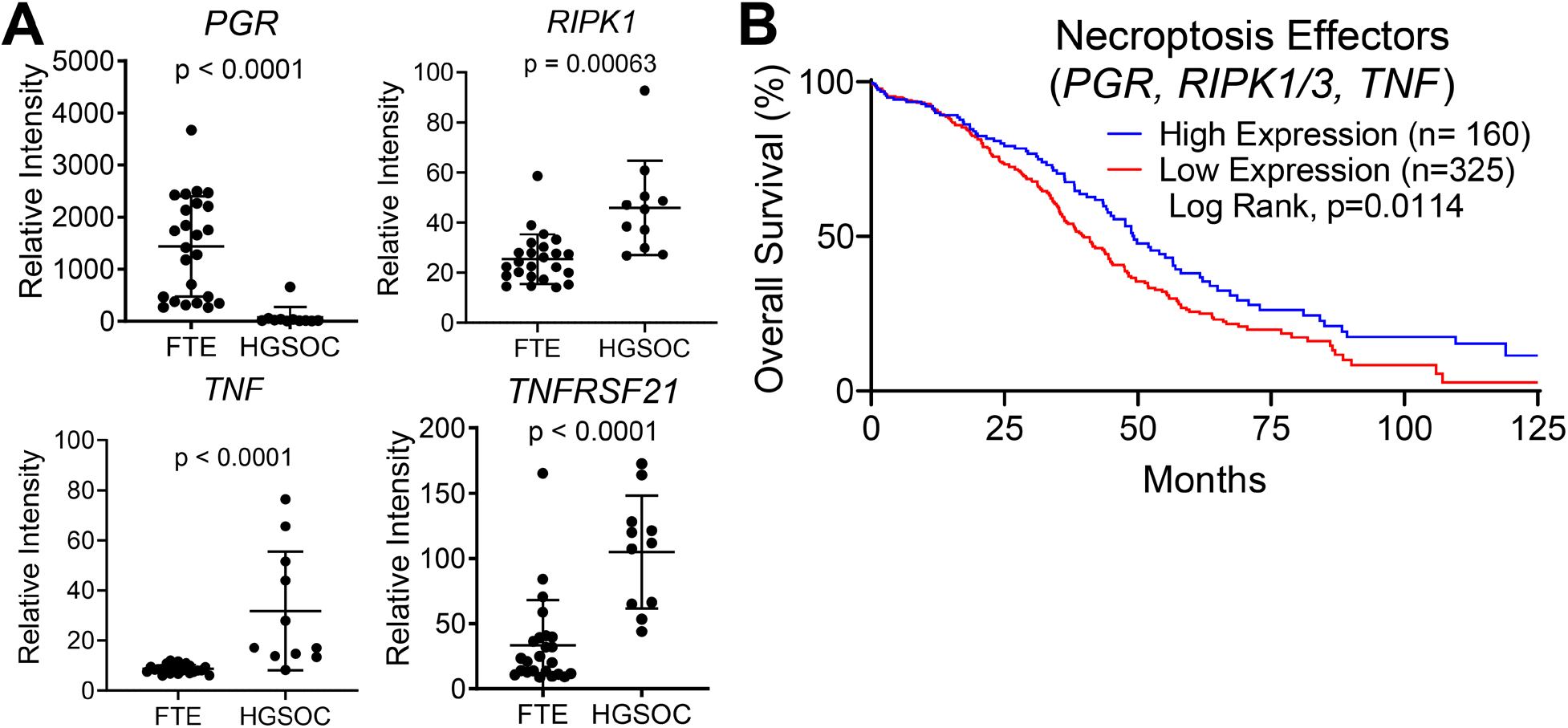
Expression of necroptosis machinery correlates positively with HGSOC patient survival. **(A)** A publicly available cohort (GSE10971) of normal FTE (n=24) and HGSOC samples (n=11) were queried for mRNA expression. Microarray data are plotted as relative intensity, including mean ± SD. P-value corrected for false-discovery rate. **(B)** A TCGA cohort of 488 HGSOC patients was queried for necroptosis effector expression. Patients were stratified into High and Low expression groups and overall survival was plotted by Kaplan-Meier analysis.

### Established HGSOC cells maintain the capacity to activate necroptosis

Wu et al. showed that P4 signaling induces necroptosis and clearance of p53-mutated FTE cells [16]. However, given the changes in necroptosis signaling in HGSOC compared to FTE, particularly the drastic downregulation of *PGR* **(Table 1** and **Fig. 1A)**, it is unknown if fully-transformed and established HGSOC cells maintain the capacity to respond to signals that activate necroptosis. We chose to test induction of necroptosis in OVSAHO cells using P4, TNFα, or (9*Z*,11*E*,13*E*)-octadecatrienoic acid, also known as α-eleostearic acid or ESA. ESA has been demonstrated to induce necroptosis in tumor cells [27] and reduce human colon cancer growth in mice [28, 29]. OVSAHO cells are PGR positive, *TP53*-mutant, have a homozygous *BRCA2* deletion, and are considered highly-representative of HGSOC tumors [30]. Following treatment with P4, TNFα, or ESA, OVSAHO cells were stained with acridine orange to distinguish cells undergoing death by necroptosis [31]. Detection was by fluorescence microscopy, and representative images are shown in **Fig. 2A**. Compared to control, all treatments increased the percentage of cells positive for necroptosis staining **(Fig. 2B)**, demonstrating that OVSAHO cells respond to P4-dependent and -independent signals to activate necroptosis. OVSAHO cells also upregulated necroptosis gene expression in response to stimulation with P4 or TNFα, including *TNF*, *RIPK1*, and *OTUD3* mRNA **(Fig. 2C)**, as well as OTUD3 protein **(Fig. 2D)**.

**Fig. 2.**
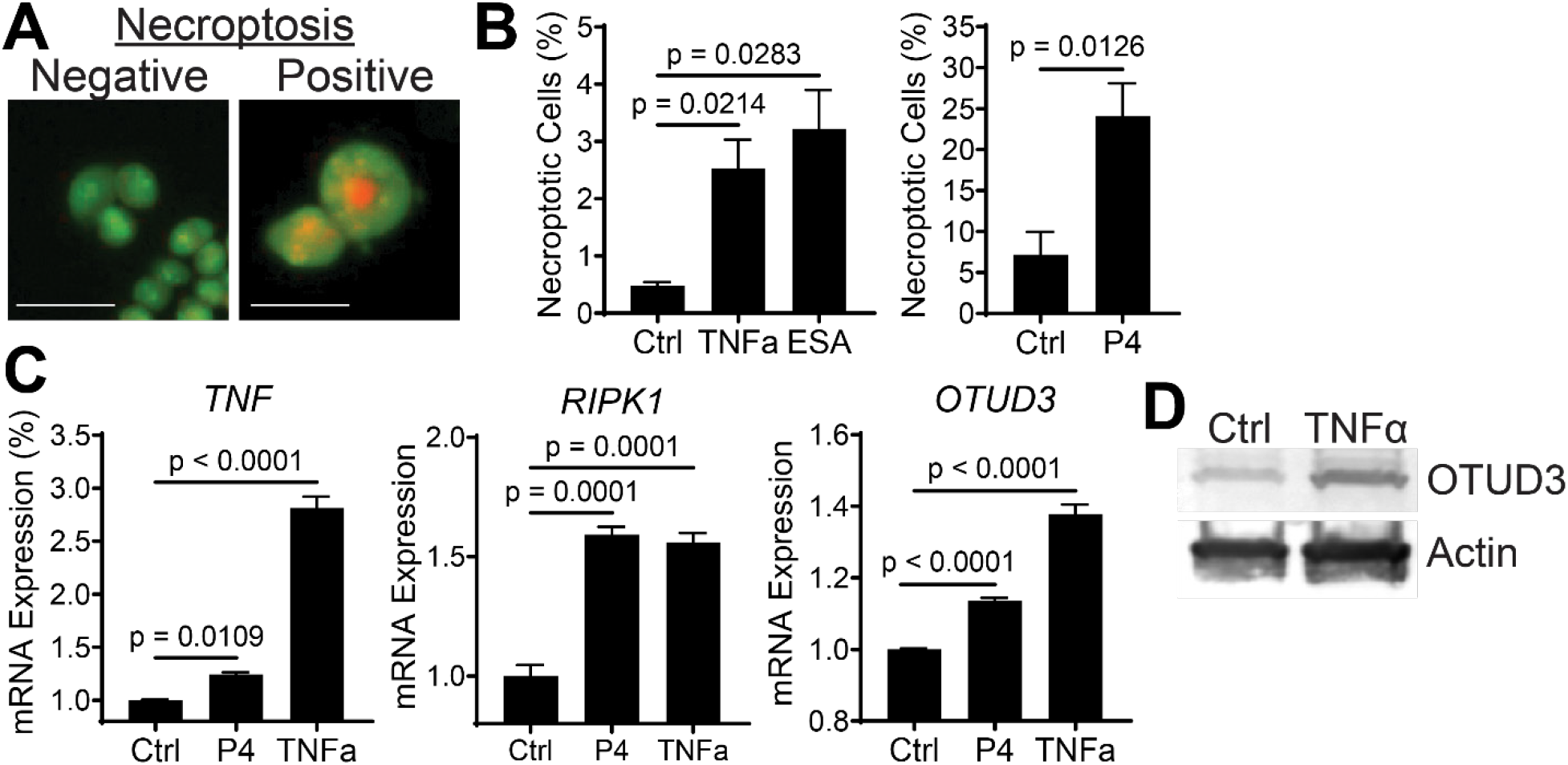
HGSOC cells activate necroptosis. **(A)** Examples of acridine orange staining to identify cells dying of necroptosis. Green = nuclei; Red/Orange = acridine orange. **(B)** OVSAHO cells were treated with DMSO control, TNFα (500 ng/mL), ESA (20 μM) or P4 (100 nM). 24 h after treatment, cells were stained with acridine orange and the percentage of positive cells was quantified. Data are plotted as mean ± SD of at least 200 cells. P-value by ANOVA or t-test. **(C)** OVSAHO cells were treated with DMSO control, P4 (10 μM), or TNFα (100 ng/mL). RT-qPCR was performed to assess mRNA expression of *TNF* and *RIPK1* at 24 h, and *OTUD3* at 48 h. Expression is shown relative to shCtrl and graphed as mean ± SD of three RT-qPCR reactions. P-value by ANOVA. **(D)** OVSAHO cells were treated with TNFα (100 ng/mL) for 48 h and OTUD3 protein expression was assayed by Western blot with β-actin as loading control.

### OTUD3 knockdown reduces PTEN and RIPK1 protein levels in HGSOC cells

Execution of necroptosis depends on post-translational modifications. For example, OTUD7B stabilizes RIPK1 through hydroxylation of lysine 11 (K11) ubiquitin chains [20], promoting RIPK1 cytosolic localization, RIPK1-RIPK3 interaction, MLKL activation, and subsequent necrotic cell death [19, 20]. OTUD3 is also a cytosolic deubiquitinating enzyme and a putative tumor suppressor. It is known to deubiquitinate and stabilize PTEN [32], but little is known about the regulation of OTUD3 or its other downstream targets. *OTUD3* expression was not significantly changed in HGSOC vs FTE in the GSE10971 cohort, but we found that higher expression of *OTUD3*, along with other necroptosis machinery, correlated with greater survival in HGSOC patients **(Fig. 1B)**. Using shRNA, we generated stable knockdowns of OTUD3 in OVSAHO cells **(Fig. 3A)** and then examined protein expression of PTEN and necroptosis effector RIPK1. A 50% knockdown of OTUD3 resulted in moderate loss of PTEN and a striking, nearly complete loss of RIPK1 protein **(Fig 3B)**.

**Fig. 3.**
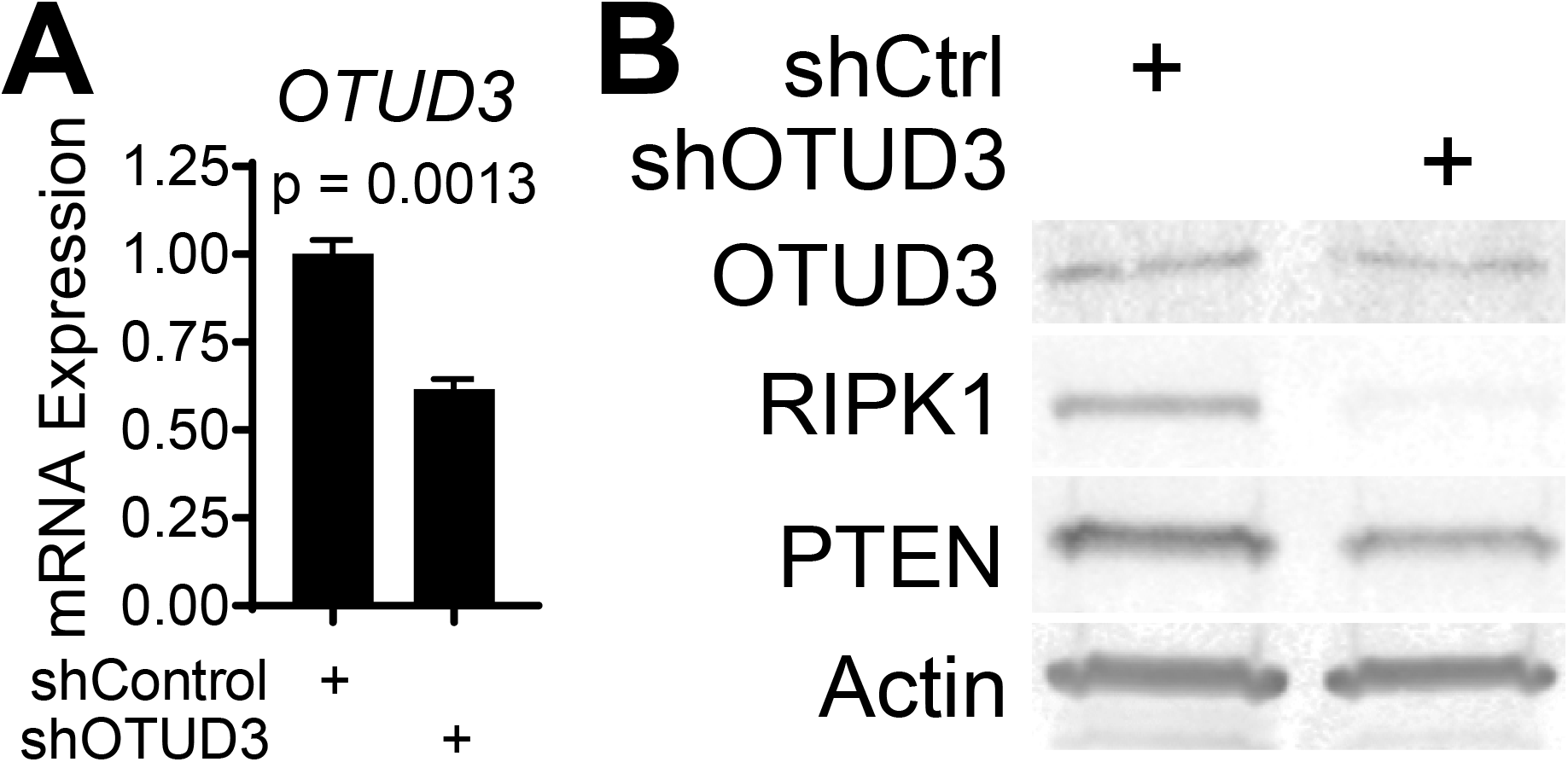
OTUD3 knockdown reduces PTEN and RIPK1 protein levels in HGSOC cells. **(A)** OVSAHO cells were stably transduced with lentivirus encoding *OTUD3* shRNA or a non-targeting control. Cells were analyzed by RT-qPCR for mRNA expression of *OTUD3* and normalized to *B2M* as internal control. Expression is shown relative to shCtrl and graphed as mean ± SD of three RT-qPCR reactions. P-value by t-test. **(B)** Control and OTUD3 knockdown cells were analyzed by Western blotting for protein expression of OTUD3, RIPK1, PTEN, and β-actin loading control.

### α-eleostearic acid reduces survival of transformed FTE and HGSOC cells

Prior to its identification as a necroptosis-inducer, ESA was shown to reduce growth of a human colon cancer in mice [28, 29]. Another report showed that ESA induces RIPK1-mediated necroptosis of tumor cells *via* generation of ROS and ATP reduction without detectable effects on normal cells [27]. Our finding that ESA induced necroptosis in OVSAHO cells **(Fig. 2B)** is aligned with those reports and suggests that pharmacologic interventions other than P4 and TNFα may be valid strategies to induce necroptosis in transformed FTE or HGSOC. Because *in vivo* use of TNFα would likely induce inflammation and other undesirable side effects [33], we chose to further examine if ESA reduces FTE or HGSOC viability. ESA treatment reduced cell viability of transformed FTE cells, as well as OVSAHO cells (HGSOC, *TP53*-mutant, *BRCA2*-deletion, PGR positive), PEO1 cells (HGSOC, *TP53*-mutant, *BRCA2*-mutant, PGR negative), and OVCAR5 (likely a gastrointestinal carcinoma [34], *TP53*-mutant*, BRCA*-wildtype, PGR status unknown) **(Fig. 4A)**. To determine if necroptosis had been induced, we stained ESA-treated cells with acridine orange [31]. PEO1 cells showed a dose-dependent response to ESA treatment **(Fig. 4B)**. RIPK1 protein expression was elevated in transformed FTE cells treated with ESA, suggesting that necroptosis activation contributed to loss of cell viability **(Fig. 4C)**. We then treated PEO1 cells with ESA and a necroptosis inhibitor, necrostatin-1 (Nec-1). Nec-1 treatment rescued cell survival of ESA-treated cells **(Fig. 4D)** and reduced acridine orange staining **(Fig. 4E)**, giving further evidence that loss of cell viability was due to activation of necroptosis.

**Fig. 4.**
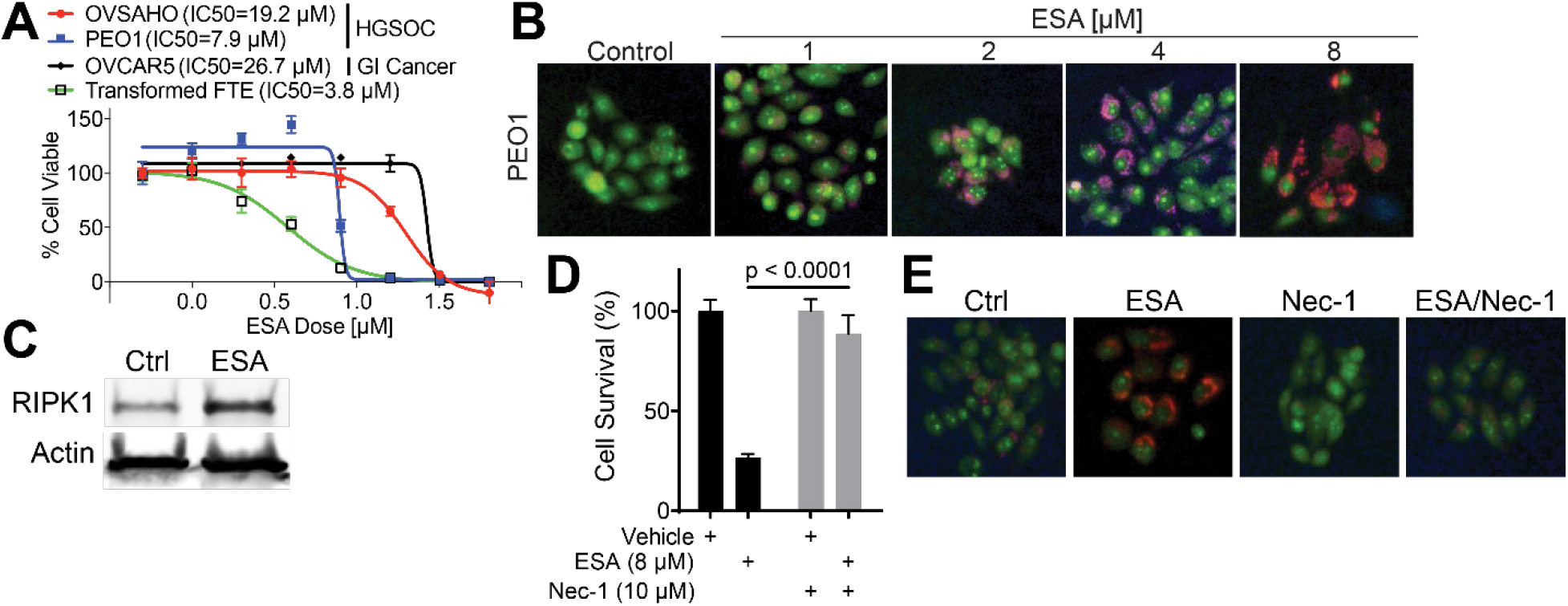
α-eleostearic acid reduces survival of transformed FTE and HGSOC cells. **(A)** Transformed FTE and cancer cells were treated with increasing doses of ESA. After 72 h, cell viability was determined by MTT assay. Data points are plotted as mean ± SD of 6 wells. Dose curves and IC50 were calculated in GraphPad Prism 8 using the sigmoidal response function. **(B)** PEO1 cells were treated with ESA or vehicle control as shown. After 24 h, cells were stained with acridine orange to identify cells undergoing death by necroptosis. Green = nuclei. Red = acridine orange. **(C)** PEO1 cells were treated with 8 μM ESA or vehicle control. After 24 h, cells were assayed by Western blot for RIPK1 protein expression and β-actin loading control. **(D)** PEO1 cells were treated with vehicle control, 8 μM ESA, and/or 10 μM Nec-1 as shown. After 48 h, the percentage of surviving cells was examined by MTT assay. Data points are plotted as mean ± SD of 6 wells. P-value by ANOVA. **(E)** PEO1 cells were treated as in **D**. After 24 hours, cells were stained with acridine orange to identify cells undergoing death by necroptosis.

## Discussion

Over 75% of HGSOC cases are diagnosed at International Federation of Gynecology and Obstetrics (FIGO) Stage III or IV, in which tumor cells have disseminated to distant tissues within the abdominopelvic cavity or have metastasized to organs beyond the peritoneum. Survival rates for such advanced disease are low, and prognosis is especially poor for postmenopausal women who have experienced a decrease in hormones including estrogen and progesterone [35]. The fact that localized, early stage HGSOC has much higher cure rates highlights that novel detection and prevention methods, as well as methods of killing tumor cells prior to extensive dissemination, will have an enormous impact on increasing survival.

The onset of HGSOC greatly increases at the time of menopause when ovarian steroid hormone production, including that of P4, greatly diminishes. DMPA significantly reduced risk of HGSOC in a large cohort, especially when duration of use was greater than three years [15], suggesting that early and/or sustained treatment may remove transformed cells. This idea is strengthened by a study showing that P4 signaling induced necroptosis and clearance of p53-mutated FTE cells [16]. However, PGR expression is frequently lost in HGSOC, a fact highlighted by our finding that *PGR* mRNA was sharply downregulated in a cohort of HGSOC when compared to nonmalignant FTE **(Table 1** and **Fig. 1A)**. Thus, while early-stage transformed FTE cells could potentially be eliminated through P4-dependent induction of apoptosis, it remains an open question whether such a mechanism could be exploited to eliminate HGSOC. High-dose megestrol did not induce an anti-tumor response in a smaller cohort of late-stage HGSOC patients [14], suggesting that targeting downstream effectors of necroptosis may be required for elimination of late-stage disease.

Despite the loss of *PGR*, numerous genes involved in TNF signaling, PGR signaling, and necroptosis were upregulated HGSOC relative to FTE, including *TNF*, *RIPK1*, and *TNFRSF21* **(Table 1** and **Fig. 1A)**. Necroptosis may be induced by P4-independent signals such as TNFα, and patients with higher levels of necroptosis machinery *PGR*, *RIPK1/3*, and *TNF* exhibit better overall survival **(Fig. 1B)**. We therefore examined necroptosis induction by P4-dependent and -independent means in HGSOC cell lines to determine if these are viable mechanisms to eliminate HGSOC cells.

Consistent with induction of necroptosis in transformed FTE [16], we observed that P4, TNFα, and ESA induced necroptosis in PGR-positive HGSOC cells (OVSAHO) **(Fig. 2A-B)**. TNFα was especially effective at inducing expression of necroptosis genes *TNF*, *RIPK1*, and *OTUD3*, as well as OTUD3 protein **(Fig 2C-D)**, suggesting that P4-independent signaling is a valid method to activate necroptosis in HGSOC cells. Consistent with the known role of OTUD3 in PTEN stability [32], knockdown of *OTUD3* decreased PTEN protein levels in OVSAHO cells. *OTUD3* knockdown also sharply decreased RIPK1 protein levels, indicating OTUD3 may function similarly to OTUD7B and that OTUD3 deubiquitinase activity [36] may regulate RIPK1 stability and subsequent necroptosis **(Fig. 3B)**. Additionally, ESA effectively induced necroptosis **(Fig. 4B)** and increased RIPK1 protein expression **(Fig. 4C)** through a P4-independent mechanism in PGR-negative HGSOC cells (PEO1). ESA response was ablated by RIPK1 inhibitor Nec-1, indicating a RIPK1-dependent activation of necroptosis **(Fig. 4D)**. Altogether, these findings indicate that HGSOC cells maintain responsiveness to necroptosis stimuli, that OTUD3 is a likely posttranslational regulator of RIPK1, and that P4 and ESA have potential therapeutic uses against transformed FTE and HGSOC cells. This is of particular interest since these compounds could be used to eliminate tumor cells while avoiding adverse side effects of TNFα [33].

Our findings suggest roles for OTUD3- and RIPK1/3-dependent necroptosis in killing HGSOC cells. However, additional key substrates of RIPK1/3 such as MLKL and subsequent ROS production have not been examined in this context. The small molecule drug necrosulfonamide has been shown to prevent an MLKL-RIPK1-RIPK3 complex from interacting with downstream necroptosis effectors [37] and may be a useful compound to further elucidate mechanisms of necroptosis activation in HGSOC. Other questions such as the roles of specific TNF receptors remain open. We identified upregulation of *TNFRSF21* in HGSOC relative to FTE **(Table 1** and **Fig. 1A)**. This finding is consistent with a report that TNFRSF21 protein is upregulated in HGSOC and that a secreted form may be a serum biomarker for HGSOC [38]. TNFRSF21 is also known as death receptor 6 (DR6) and signaling through DR6 is known to induce necroptosis in endothelial cells [39]. It is unknown if signaling through DR6 can be exploited to specifically eliminate high-DR6-expressing HGSOC cells without severe side effects against endothelial cells.

## Conclusions

Our findings further highlight the potential of exploiting necroptosis activation to eliminate HGSOC or as a preventative measure to eliminate precursor transformed FTE cells. Future *in vitro* studies will elucidate exact necroptosis activation mechanisms and dependencies, while *in vivo* studies targeting specific effectors such as DR6, OTUD3, RIPK1/3, and MLKL will determine if necroptosis can be exploited to effectively kill tumor cells while avoiding adverse side effects.

## Methods

### Ovarian cancer dataset analysis

Gene expression was compared between nonmalignant FTE and HGSOC groups in a publicly available cohort (GSE10971). Genes for evaluation were selected using gene ontology (necroptosis, ID 0070266; PGR signaling, ID 0050847; TNF signaling, 0010803), as well as *BRCA1*, *BRCA2*, TNF receptors and OTUD family genes. P-values were adjusted for false-discovery rate by the Benjamini-Hochberg method [40]. An ovarian cancer TCGA cohort [11] was analyzed for mRNA expression of necroptosis effectors. Patients were stratified by High and Low expression and overall survival was plotted using Kaplan Meier analysis.

### Cell culture, shRNA, and lentivirus

Cell lines were obtained from the Gynecologic Tumor and Fluid Bank (GTFB) at the University of Colorado and were authenticated at the University of Arizona Genomics Core using short tandem repeat DNA profiling. Regular Mycoplasma testing was performed using MycoLookOut PCR (Sigma). OVSAHO, PEO1, and OVCAR5 were cultured in RPMI 1640 supplemented with 10% fetal bovine serum (FBS) and 1% penicillin/streptomycin. 293FT lentiviral packaging cells were cultured in DMEM supplemented with 10% FBS and 1% penicillin/streptomycin. All cells were grown at 37 °C supplied with 5% CO_2_. shRNA in pLKO.1 lentiviral vector plasmids were purchased from the University of Colorado Functional Genomics Facility. Lentivirus was packaged as previously described [41] in 293FT using 3^rd^ generation packaging plasmids (Virapower, Invitrogen) with polyethyleneimene (PEI) transfection in a 1:3 DNA:PEI ratio. Culture supernatant was harvested at 48-72 h post-transfection and processed through 0.45 μM filters. Viruses encoded a puromycin resistance gene. Transduced cells were selected in 1 μg/mL puromycin. OTUD3 was knocked down using TRCN000253744. An empty pLKO.1 was used as a control.

### Acridine orange staining for necroptosis

Following treatment, cells were washed 1X with PBS. Acridine orange (Sigma #113000) was added to the cells at a final concentration of 5 μg/mL in PBS and incubated for 15 min at 37 °C. Cells were washed 2X with PBS and immediately imaged on an Olympus FV1000 inverted fluorescence microscope. Images of cells were counted in ImageJ (NIH). At least 200 cells were counted per condition.

### Reverse-transcriptase quantitative PCR (RT-qPCR)

RNA was isolated from cells using the RNeasy Plus Mini Kit (Qiagen). mRNA expression was determined using SYBR green Luna One Step RT-qPCR Kit (New England BioLabs) on a C1000 Touch (Bio-Rad) or QuantStudio 6 (Applied Biosystems) thermocycler. Expression was quantified by the ΔΔCt method using target-specific and control primers. β-2-microglobulin (*B2M*) was used as internal control. mRNA-specific primers were designed to span exon-exon junctions to avoid detection of genomic DNA. Primer sequences: *TNF-F:* AGCCTCTTCTCCTTCCTGAT, *TNF-R:* CCAGAGGGCTGATTAGAGAGA; *RIPK1-F:* TCTGTGTTTCCACAGAACCC, *RIPK1-R:* GTTCATCATCTTCGCCTCCTC; *OTUD3-F:* CACCTCCCGCAGCTTCA, *OTUD3-R:* GCAGCCATGTCCCGAAAG; *B2M-F:* GGCATTCCTGAAGCTGACA, *B2M-R:* CTTCAATGTCGGATGGATGAAAC.

### Immunoblotting

For total protein, samples were lysed and briefly sonicated in RIPA buffer (150mM NaCl, 1% TritonX-100, 0.5% sodium deoxycholate, 0.1% SDS, 50mM Tris pH 8.0) supplemented with cOmplete EDTA-free protease inhibitors (Roche #11873580001) and phosphatase inhibitors NaF and NaV. Protein was separated by SDS-PAGE and transferred to PVDF membrane using the TransBlot Turbo (BioRad). All subsequent blocking and antibody incubations were performed in LI-COR Odyssey buffer (LI-COR #927-50000). Membranes were blocked for 1 hour at room temperature. Primary antibody incubation was performed overnight at 4 °C. Membranes were washed 3 times for 5 minutes each in TBST (50 mM Tris pH 7.5, 150 mM NaCl, 0.1% Tween-20), then LI-COR fluorophore-labeled secondary antibodies (goat anti-rabbit [#925-68071 or #926-32211] or goat anti-mouse [#926-68070 or #925-32210] were applied at 1:20,000 dilution for one hour at room temperature. Membranes were washed again 3 times for 5 minutes each in TBST. Bands were visualized using the LI-COR Odyssey Imaging System. Primary antibodies (catalog number, species, working concentration): β-actin (Abcam ab6276, mouse, 1:10,000); OTUD3 (Invitrogen PA5-24508, rabbit, 1:1000 or PA5-55895, rabbit, 1:1000); RIPK1 (Cell Signaling Technology #3493, rabbit, 1:1000). PTEN (Cell Signaling Technology #9559, rabbit, 1:1000).

### Cell viability assay

10,000 cells were plated in 96-well plates and treated with ESA and/or Nec-1 or vehicle control as described in the Figure legends. After treatment, cell viability was assessed by MTT assay (Promega #G3582) using the manufacturer’s protocol.

### Software and statistical analysis

Statistical analysis of GSE10971 was performed using GEO2R functions, including adjustment of P-values for false discovery rate. For other analyses, calculation of P-value was performed using GraphPad Prism 8. Quantitative data are expressed as mean ± SD unless otherwise stated. Two-tailed t-test was used for single comparisons. Analysis of variance (ANOVA) with Fisher’s Least Significant Difference (LSD) was used in multiple comparisons. For all statistical analyses, the level of significance was set at 0.05.

## Declarations

### Ethics approval and consent to participate

Not applicable

### Consent for publication

Not applicable

### Availability of data and materials

Data generated and/or analyzed during this study are available from the corresponding author on reasonable request. Ovarian patient cohort data are publicly available.

### Author’s contributions

JJ – concept, experimental design, data analysis, manuscript preparation; ECB – experiment execution; BGB – concept, experimental design and execution, manuscript preparation; ZLW – experimental execution and manuscript preparation

## Acknowledgements

The authors thank Elizabeth Woodruff, PhD for assistance with manuscript preparation and editing.

